# BUB1 inhibition sensitizes lung cancer cell lines to radiotherapy and chemoradiotherapy

**DOI:** 10.1101/2024.04.19.590355

**Authors:** Shivani Thoidingjam, Sushmitha Sriramulu, Oudai Hassan, Stephen L. Brown, Farzan Siddiqui, Benjamin Movsas, Shirish Gadgeel, Shyam Nyati

## Abstract

**Background:** Lung cancer is a major public health concern, with high incidence and mortality. Despite advances in targeted therapy and immunotherapy, microtubule stabilizers (paclitaxel, docetaxel), DNA intercalating platinum drugs (cisplatin) and radiation therapy continue to play a critical role in the management of locally advanced and metastatic lung cancer. Novel molecular targets would provide opportunities for improving the efficacies of radiotherapy and chemotherapy.

**Hypothesis:** We hypothesize that BUB1 (Ser/Thr kinase) is over-expressed in lung cancers and that its inhibition will sensitize lung cancers to chemoradiation.

**Methods:** BUB1 inhibitor (BAY1816032) was combined with platinum (cisplatin), microtubule poison (paclitaxel), a PARP inhibitor (olaparib) and radiation in cell proliferation and radiation sensitization assays. Biochemical and molecular assays were used to evaluate their impact on DNA damage signaling and cell death mechanisms.

**Results:** BUB1 expression assessed by immunostaining of lung tumor microarrays (TMAs) confirmed higher BUB1 expression in NSCLC and SCLC compared to that of normal tissues. BUB1 overexpression in lung cancer tissues correlated directly with expression of TP53 mutations in non-small cell lung cancer (NSCLC). Elevated BUB1 levels correlated with poorer overall survival in NSCLC and small cell lung cancer (SCLC) patients. A BUB1 inhibitor (BAY1816032) synergistically sensitized lung cancer cell lines to paclitaxel and olaparib. Additionally, BAY1816032 enhanced cell killing by radiation in both NSCLC and SCLC. Molecular changes following BUB1 inhibition suggest a shift towards pro-apoptotic and anti-proliferative states, indicated by altered expression of BAX, BCL2, PCNA, and Caspases 9 and 3.

**Conclusion:** A direct correlation between BUB1 protein expression and overall survival was shown. BUB1 inhibition sensitized both NSCLC and SCLC to various chemotherapies (cisplatin, paclitaxel) and targeted therapy (PARPi). Furthermore, we present the novel finding that BUB1 inhibition sensitized both NSCLC and SCLC to radiotherapy and chemoradiation. Our results demonstrate BUB1 inhibition as a promising strategy to sensitize lung cancers to radiation and chemoradiation therapies.

## 1. Introduction

Lung cancer is the leading cause of cancer-related deaths worldwide and approximately a quarter million new cases and half that number of deaths are expected in 2024 in the United States alone (1). Lung cancer is categorized into two primary subtypes: NSCLC and SCLC. While NSCLC includes histological subtypes such as lung adenocarcinoma (LUAD), lung squamous cell carcinoma (LUSC), and large cell carcinoma (LCLC), constituting 85% of cases, SCLC accounts for the remaining 15% (2). Despite advancements in treatment modalities, both subtypes pose significant treatment challenges, including late-stage diagnoses and resistance to various therapies (2). The average 5-year survival rate for lung cancer is 25% (1). Checkpoint blockade has become popular in NSCLC treatment (3), and ongoing trials are assessing targeted agents and checkpoint inhibitors in adjuvant therapy for early-stage NSCLC (4, 5). While molecularly targeted therapies and immune checkpoint inhibitors improve the outcome for NSCLC, a substantial proportion of NSCLC patients don’t derive significant clinical benefits from these therapies (6). Multimodality treatment approach incorporating radiation and targeted therapy for management of lung cancer is popular (7). There is thus an urgent need to identify novel therapeutic targets that can improve outcome and overcome therapeutic resistance.

BUB1 is a highly conserved serine/threonine kinase and plays a critical role in spindle assembly checkpoint (SAC) function, facilitating congression in metaphase, and safeguarding sister chromatid cohesion (8). BUB1 also plays a crucial role in DNA damage response (9, 10). Dysregulated BUB1 expression is implicated in various malignancies, including lung cancers (11–13). Our previous studies established BUB1’s kinase activity driving aggressive cancer phenotypes via TGF-β signaling (14, 15). Pharmacological inhibition of BUB1 significantly reduced tumor xenograft growth (12, 16) and sensitized cancer cells to taxanes, ATR, or PARP inhibitors confirming BUB1 as a therapeutic target for enhancing the efficacy of chemotherapies and targeted therapies (17). These studies, however, did not evaluate arole for BUB1 in improving the efficacies of radiotherapy or chemoradiation in lung cancers, especially SCLC. In this study we demonstrate that BUB1 inhibition sensitizes models of NSCLC and SCLC to radiotherapy (BUB1i + radiotherapy) and chemoradiation (BUB1i + chemo/targeted therapy + radiation). Furthermore, a rationale is provided for designing novel combination clinical trials targeting BUB1 along with standard of care therapies.

## 2. Materials and Methods

### 2.1. BUB1 expression in lung cancer

BUB1 expression was assessed using the Lung Cancer Explorer (LCE) (18) and UALCAN (19) databases. The TCGA_LUAD_2016, TCGA_LUSC_2016, and Rousseaux_2016 datasets from LCE (18) were used to generate BUB1 expression and survival data.

### 2.2. Immunohistological analysis of BUB1 expression in lung tumor microarrays

Formalin-fixed, paraffin-embedded NSCLC and SCLC TMAs were procured from Tissue array (https://www.tissuearray.com). The TMA’s (LUC1201MSur 120 cases and SCLC681Sur 68 cases) were immunostained with an IHC-verified BUB1 antibody (1:50 dilution; Abcam #ab195268). Staining intensity and distribution were scored by a pathologist who was blind to the clinical parameters.

### 2.3. Cell culture

NSCLC (A549, H2030, H1975, Calu-1) and SCLC (NCI-H2198, and NCI-H1876) cell lines were obtained from ATCC. These cells carry different driver mutations (**Table 1**). Cell lines were cultured in DMEM (ATCC, 30-2002), McCoy (ATCC, 30-2002), RPMI-1640 (ATCC, 30-2001) or HITES media (ATCC, 30-2006) which was supplemented with 5-10% Fetal Bovine Serum (Thermo Fisher Scientific, FB12999102) and Penicillin-Streptomycin (5,000 U/mL) (Thermo Fisher Scientific, 15070063). Cells were maintained at 37°C with 5% CO_2_ and were regularly tested for Mycoplasma contamination using the MycoAlert PLUS kit (Lonza, Cat. No. LT07 −705).

**Table 1.**
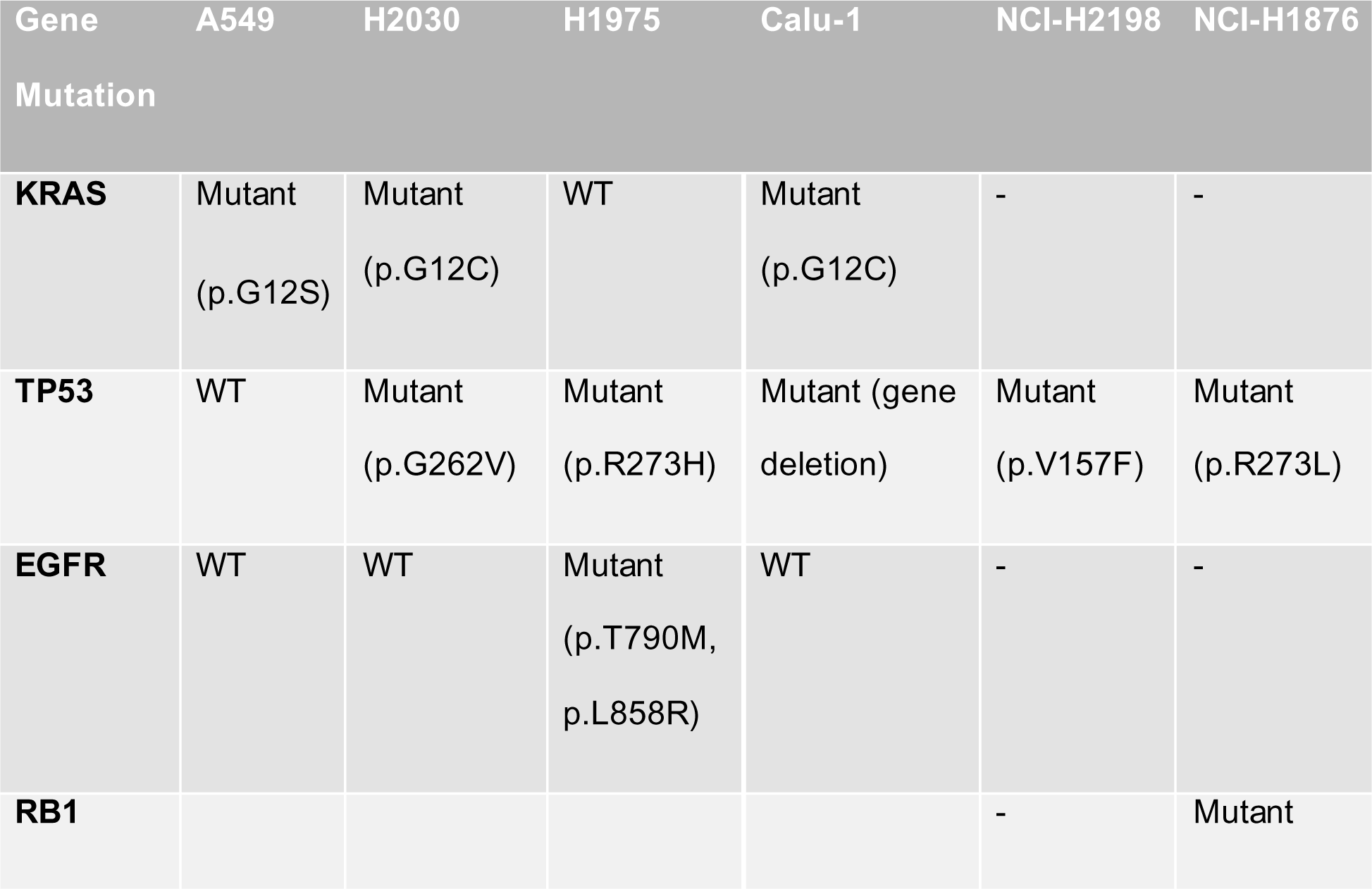
Gene mutations in lung cancer cell lines.

### 2.4. Drug treatment and Radiation

BUB1 inhibitor BAY1816032 (CT-BAY181), paclitaxel (CT-0502), and olaparib (AZD2281, CT-A2281) were obtained from Chemitek, and cisplatin (PHR1624-200MG) was sourced from Millipore Sigma. DNAPK inhibitor NU7441 (S2638) was acquired from Selleck Chemicals. Stock solutions of BAY1816032 (20 mM), paclitaxel (10mM), cisplatin (20mM), and olaparib (20mM) were made in DMSO, aliquoted and stored at −80°C. Working solutions with varying drug concentrations in culture media were freshly prepared to maintain drug stability and activity during assays. A CIX3 cabinet irradiator [Xstrahl Inc. Suwanee GA] equipped with a 320kV metal ceramic X-ray tube and operated at 10mA was used for radiation exposures. A 1 mm copper filter was used for beam hardening. Cell monolayers in plastic 6 well tissue culture plates were exposed to 2, 4, and 6 Gy radiation. Quality assurance testing included a calibrated and certified small field electrometer and routine use of x-ray sensitive film (EBT3 Gafchromic) to confirm radiation delivery.

### 2.5. Cell survival assay

Cells were seeded at 1000 cells per well in a 96-well plate. After 24 hours, cells were treated with BAY1816032, cisplatin, paclitaxel, and olaparib at concentrations ranging from nanomolar to micromolar (**Table 2**). Cells were treated for 3 days or 7 days depending on their doubling time. Cell proliferation was assessed using the AlamarBlue® Cell Viability Reagent (Thermo Fisher Scientific, Cat. No. DAL1100) according to the manufacturer’s protocol. Data were analyzed using GraphPad Prism V8, and P-values were calculated to determine the significance.

**Table 2.**
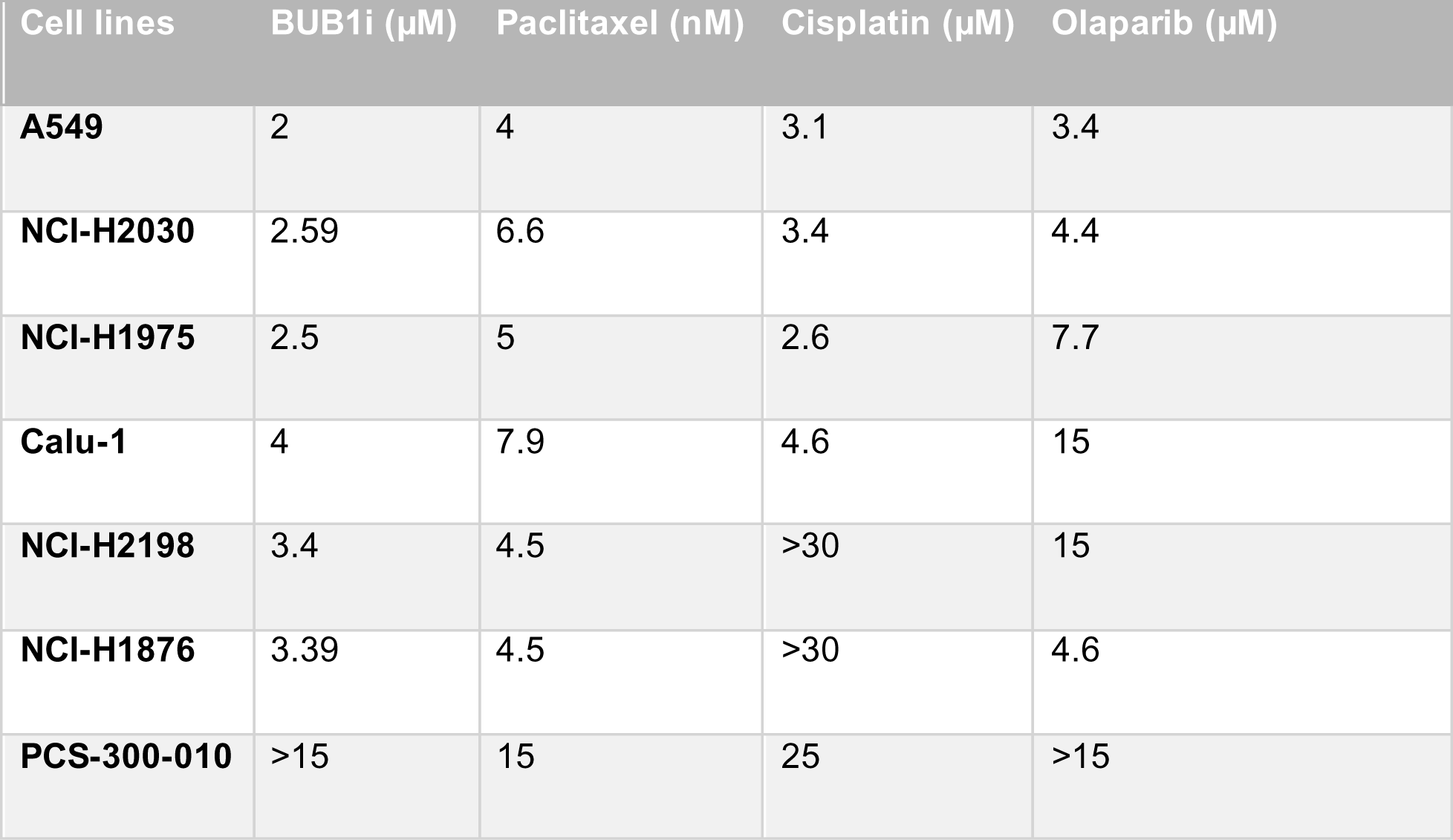
Half-maximal inhibitory concentration (IC_50_) of Chemotherapy drugs in lung cancer cell lines.

| Cell lines | BUB1i (μM) | Paclitaxel (nM) | Cisplatin (μM) | Olaparib (μM) |
| --- | --- | --- | --- | --- |
| **A549** | 2 | 4 | 3.1 | 3.4 |
| **NCI-H2030** | 2.59 | 6.6 | 3.4 | 4.4 |
| **NCI-H1975** | 2.5 | 5 | 2.6 | 7.7 |
| **Calu-1** | 4 | 7.9 | 4.6 | 15 |
| **NCI-H2198** | 3.4 | 4.5 | >30 | 15 |
| **NCI-H1876** | 3.39 | 4.5 | >30 | 4.6 |
| **PCS-300-010** | >15 | 15 | 25 | >15 |

### 2.6. Clonogenic cell survival assay

The clonogenic cell survival assay was performed to determine the radiation sensitization of BUB1 inhibitor alone or in combination with chemotherapies or targeted therapy (PARPi). Cells were seeded in 6-well tissue culture plates at different densities in triplicates. After 24 hours, cells were treated with several drug concentrations (BAY1816032, paclitaxel, cisplatin, and olaparib), alone or in combinations and irradiated (0, 2, 4, and 6 Gy) an hour after drug treatment. Cells were allowed to grow for 10-15 days until colonies with 50 or more cells formed. Colonies were fixed, stained, visually inspected, and counted to calculate plating efficiency (PE), survival fractions (SF) and radiation enhancement ratios (rER).

### 2.7. DNA damage assay

DNA damage was assessed using the γH2AX immunofluorescence assay. 1.5 x 10^5^cells were seeded on 12 mm cover-glasses in a 6-well plate and after 24 hours of attachment treated with BUB1i (1µM). After 1 hour, cells were irradiated (4 Gy) and assessed for damage at 30 minutes, 4-, 16 -, and 24-hours post-radiation. Cells were fixed, permeabilized, and blocked before incubation with anti-phospho-Histone H2A.X (Ser139) antibody (Millipore Sigma, Cat. No. 05-636-25UG) followed by Alexa Fluor 568 goat anti-mouse IgG (H+L) (Invitrogen, Cat. No. A-11004). Nuclei were counterstained with DAPI, and images were captured using a fluorescent microscope (Zeiss, Germany), with an average of 100 nuclei per image. Cells with more than 10 foci were counted as being positive for γH2AX. At least 3 random fields/samples were counted. Each experiment was repeated at least three times.

### 2.8. Quantitative real-time PCR

For Quantitative real-time polymerase chain reaction (qPCR) studies, 1 x 10^5^ cells were seeded in 6-well plates. After 24 hours, cells were treated with 1 μM BUB1i or DNAPKi for 72 hours. Total RNA was extracted using Trizol reagent (Thermo Fisher Scientific, 15596026) and RNA concentration and purity were assessed using Nanodrop (Thermo Fisher Scientific, Nanodrop 2000c). First-strand cDNA synthesis was performed using SuperScript III Reverse Transcriptase (Thermo Fisher Scientific, Cat. No. 18080044), dNTPs (Thermo Fisher Scientific, Cat. No. R0191), and random primers (Thermo Fisher Scientific, Cat. No. 48190011). qPCR reactions were conducted in triplicate for each sample using Takyon Low ROX SYBR 2X MasterMix blue dTTP (Eurogentec, Cat. No. UF-LSMT-B0701) and KiCqStart® SYBR® Green pre-designed primers (**Table 3**) on a QuantStudio 6 Flex Real-Time PCR system. GAPDH gene served as a sample control (**Table 3**). Comparative Ct method (or ΔCt) was used to analyze the data.

**Table 3.**
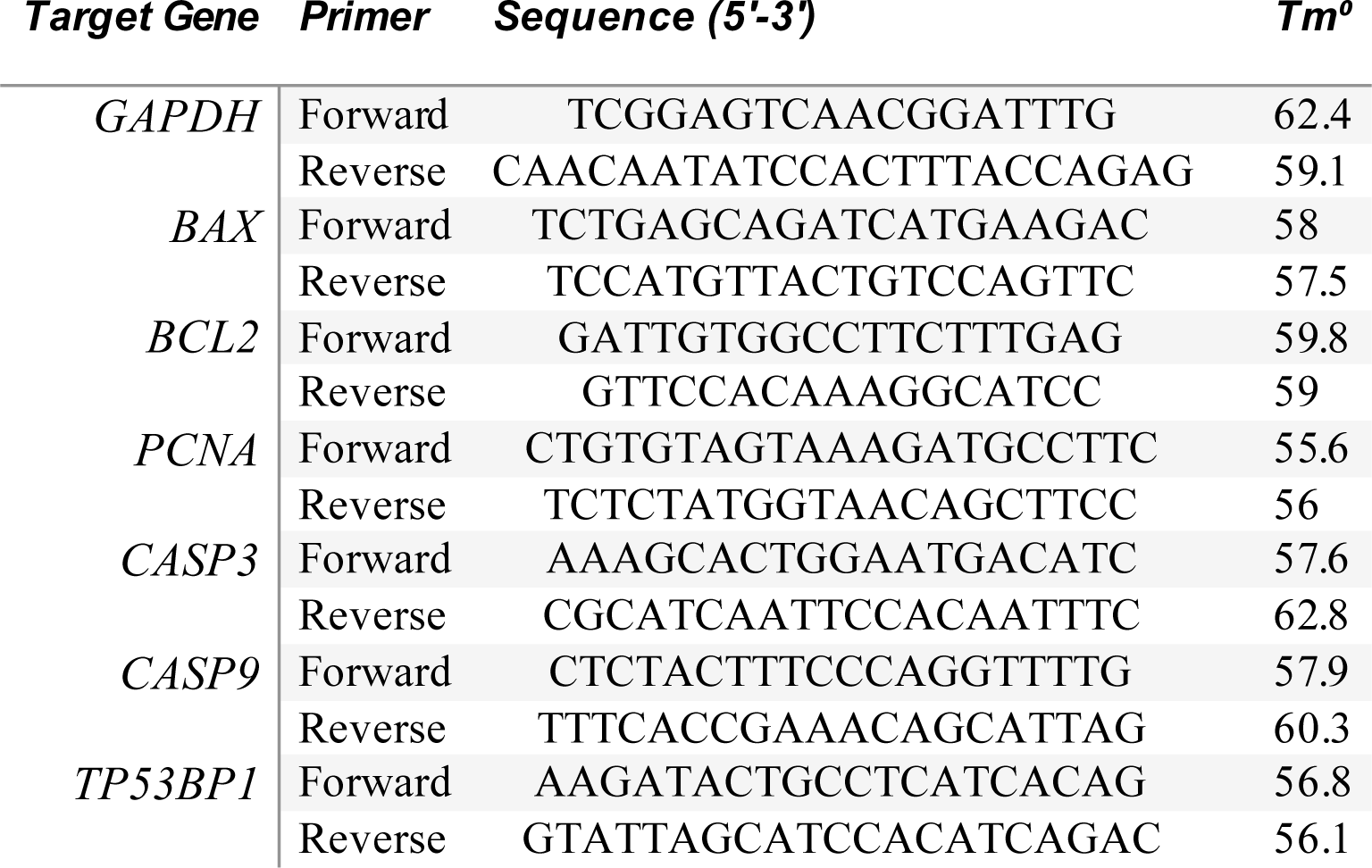
Primer sequences used in quantitative PCR (qPCR)

### 2.9. Statistical analysis

BUB1 expression data were analyzed using unpaired t-tests and ordinary one-way ANOVA. Survival analyses were conducted using the Kaplan-Meier method. Cell proliferation data was analyzed using a nonlinear regression curve fit. The Combinatory Index (C.I.) was determined using Compusyn software, and the Chow-Talay C.I. calculation formula: C.I. = (D)1 /(Dχ)1 + (D)2/(Dχ)2, where (Dχ)1 and (Dχ)2 represent the concentrations of each drug alone to achieve χ% effect, while (D)1 and (D)2 denote the concentrations of drugs in combination to produce the same effect. A C.I. value less than 1, equal to 1, and greater than 1 indicated synergism, additivity, and antagonism, respectively, as described previously (20). qPCR data was analyzed by ordinary one-way ANOVA with Dunnett’s multiple comparisons test. Graphs were generated using GraphPad Prism v8. Results from three independent experiments were presented as mean ± standard error of the mean (SEMs), with p-values of 0.05 or less considered statistically significant.

## 3. Results

### 3.1. BUB1 is overexpressed in LUAD, LUSC and SCLC, and is correlated with poorer survival

BUB1 was significantly overexpressed in LUAD (n=517, p-val=7.4e-40) (**Figure 1A**) and LUSC (n=501, p-val=8.1e-34) (**Figure 1B**) compared to normal tissues (n=59, 51) from TCGA data. BUB1 was expressed higher in LUSC (n = 100, p-val=1e-17and SCLC (n = 21, p-val=1.5e-10) as compared to LUAD (n = 85) (**Figures 1C-D**). Higher BUB1 expression correlated with inferior overall survival in LUAD patients (p-val = 0.0033 (**Figure 1E**). Additionally, BUB1 was expressed at higher levels in TP53 mutated LUAD (n = 233) and LUSC (n = 369) compared to TP53 non-mutant cases (UALCAN TCGA database) (**Figures 1F-G**), suggesting a potential association or correlation between TP53 mutation status and BUB1 expression in these specific lung cancer subtypes.

**Figure 1:**
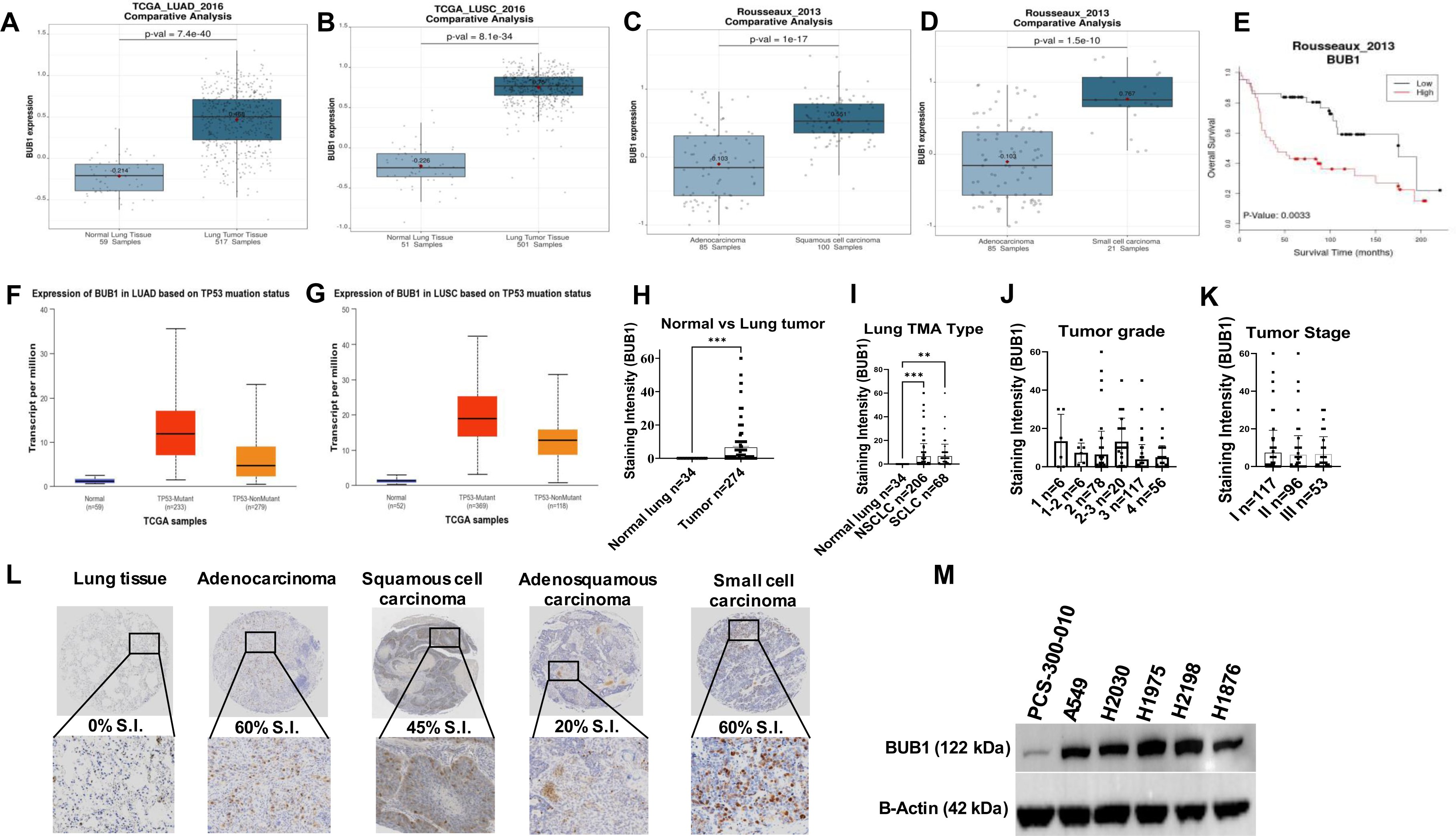
(A) BUB1 is overexpressed in LUAD (n = 517, p-val=7.4e-40) compared to normal tissues (n = 59) (Lung explorer, TCGA_LUAD_2016 database). (B) BUB1 is overexpressed in LUSC (n = 501, p-val=8.1e-34) compared to normal tissues (n = 51) (TCGA_LUSC_2016 database). (C) & (D) BUB1 expression levels are in LUSC (n = 100, p-val=1e-17) and SCLC (n = 21, p-val=1.5e-10) are higher than LUAD (n = 85) Rousseaux_2013 database). High BUB1 expression correlates with poorer survival in (E) LUAD, p-val=0.0033 compared to low BUB1 (Rousseaux_2013 database). (F) & (G) Higher BUB1 expression is seen in TP3 mutant LUAD (n = 233) and LUSC (n = 369) (UALCAN TCGA database). TMA analysis: (H) Higher BUB1 expression is seen in lung tumor samples compared to normal tissues. (I) NSCLC (n = 206) and SCLC (n = 68) exhibited higher BUB1 expression compared to normal tissues (n =34). (J) & (K) No correlation between BUB1 expression and tumor grade and stage was seen in the TMAs analyzed. (L) BUB1 expression in TMA of different types of lung cancers with staining intensity. (M) BUB1 basal level expression in lung cancer cell lines.

### 3.2. BUB1 protein is overexpressed in tumor tissues

BUB1 immunostaining in lung TMAs revealed BUB overexpression in tumor tissues (n=274) compared to normal tissues (n=34) (**Figure 1H**). Within these tumors, BUB1 was overexpressed in NSCLC (n=206) and SCLC (n=68) compared to normal lung tissues (n=34) (**Figure 1I)**. However, we did not observe any significant correlation between BUB1 expression and tumor grades or stages (**Figures 1J-K)**. Representative images demonstrating differential BUB1 immunoreactivity (**Figure 1L**). BUB1 is expressed at very low levels in normal lung epithelial cell line PCS-300-010 compared to NSCLC and SCLC cell lines (**Figure 1M**). We observed highest BUB1 expression in NCI-H2198 and NCI-H1975, followed by A549, NCI-H2030, and NCI-H1876.

### 3.3. BUB1 inhibition suppresses cell proliferation and clonogenic capacity

The BUB1 inhibitor, BAY1816032, dose-dependently suppressed proliferation of lung cancer cells. The IC_50_ ranged from 2-4μM in A549, H2030, H1975, Calu-1, NCI-H2198, and NCI-H1876 cells using the Alamar Blue cytotoxicity assay (**Figures 2B-G**). The IC_50_ was found to be substantially higher (15μM), that is less toxic, to normal lung epithelial PCS-300-010 cells (**Figure 2A**). The IC_50_ for single-agent BUB1i dropped to nano-molar range in clonogenic survival assays in A549, H2030, and H1975 cells (500 - 900 nM; **Figures 2I-K**) indicating that in multiple lung cancer cell lines, BUB1 inhibition is highly toxic. The SCLC cell lines did not form well-differentiated colonies in the culture conditions used, thus the effect of BAY1816032 in these cells could not be evaluated by clonogenic survival assay.

**Figure 2:**
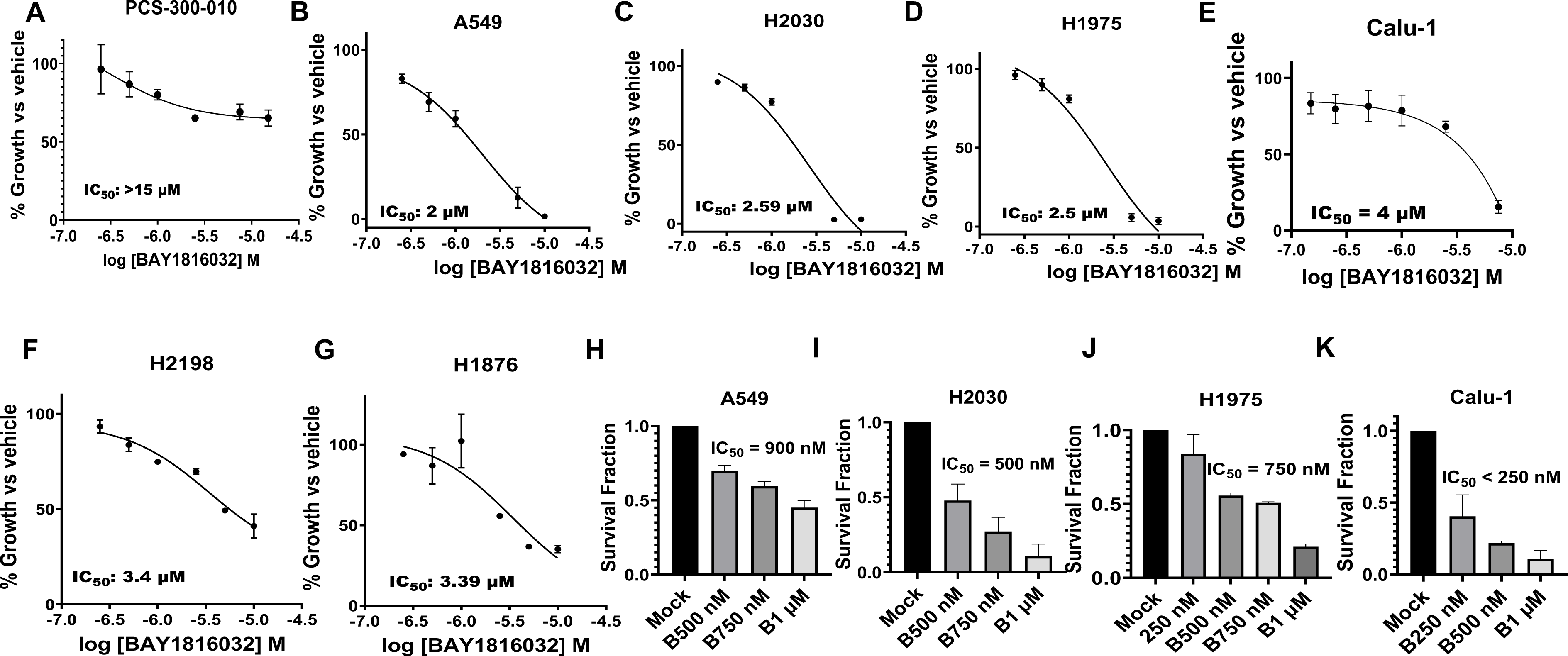
BUB1 inhibition causes cell death/toxicity in Alamar blue cell proliferation assay in normal lung (A) PCS-300-010 and lung cancer cells (B) A549 (C) H2030 (D) H1975 (E) Calu-1 (F) H2198 and (G) H1876. Clonogenic Survival Assays plots for (H) A549 (I) H2030 (J) H1975 (K) Calu-1 shown in the bar graphs.

### 3.4. The sequence and doses of BUB1 inhibitor determine sensitivity to paclitaxel, cisplatin and olaparib

NSCLC and SCLC cell lines displayed varying toxicities to single agent paclitaxel, cisplatin, and olaparib, with IC_50_ values ranging from 4 to 7.9 nM for paclitaxel, 2.6 to >7.7 μM for cisplatin, and 3.4 to >15 μM for olaparib (**Figure 3** and **Table 2)**. Different treatment strategies were explored in NCI-H1975 cells including single agent (**Figures 4A-C**), sequential and concurrent treatments (**Figures 4D-O**). Pretreatment with BUB1i 24 hours prior to cisplatin resulted in antagonism (C.I. = 1.22 **Figure 4D**), further potentiated by raising cisplatin concentration with constant BUB1i dose (C.I. = 1.27; **Figure 4D**). Conversely, pretreatment with cisplatin followed by BUB1i exhibited antagonism (C.I. = 1.5, **Figure 4E**), alleviated by increasing BUB1i dose while maintaining constant cisplatin dose (C.I. = 1.1; **Figure 4E**). In concurrent BUB1i and cisplatin treatment, increasing cisplatin concentration with a constant BUB1i dose shifted from synergism to additivity (C.I. = 0.9 to 1.1; **Figure 4F**), and increasing BUB1i dose with constant cisplatin switched antagonism to additivity (C.I. = 1.5 to 1.1; **Figure 4G**). For BUB1i and paclitaxel, sequential treatment displayed additive effects (C.I. >1; **Figures 4H-I**), while concurrent treatment showed synergism (C.I. <1; **Figures 4J-K**). In the sequential BUB1i and olaparib combination, BUB1i pretreatment exhibited additive effects (**Figure 4L)**, while olaparib pretreatment demonstrated synergistic effects (**Figure 4M**). In concurrent BUB1i and olaparib treatment, survival fractions were very low (lower than sequential treatment) and thus C.I. could not be calculated **(Figures 4N-O)**. Based on these results, concurrent treatment was selected as the preferred approach for drug combinations in subsequent studies.

**Figure 3:**
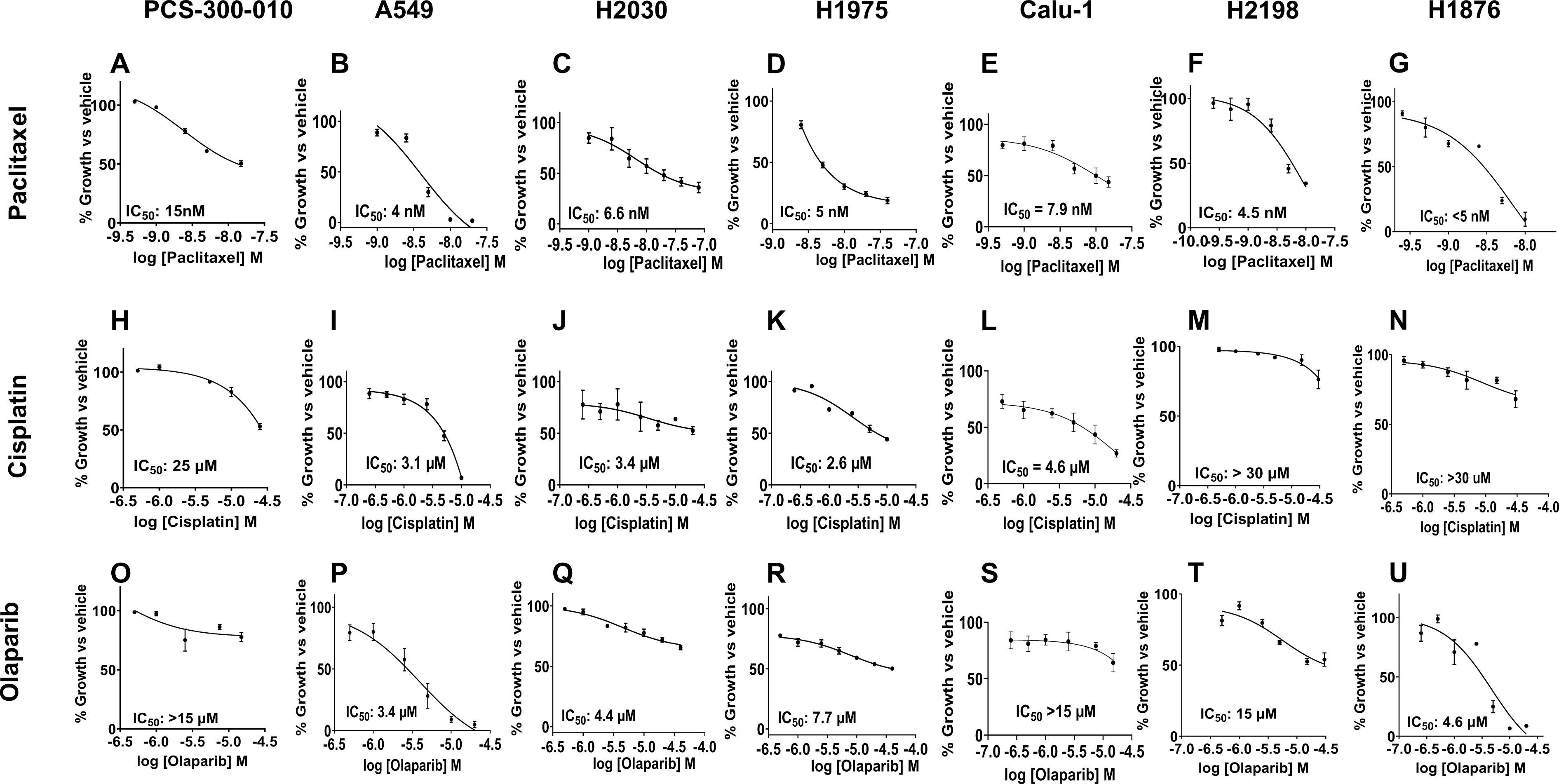
Estimation of cytotoxic concentration (IC_50_) of chemotherapeutic drugs paclitaxel, cisplatin and olaparib in lung cancer cell lines by Alamar blue assay.

**Figure 4:**
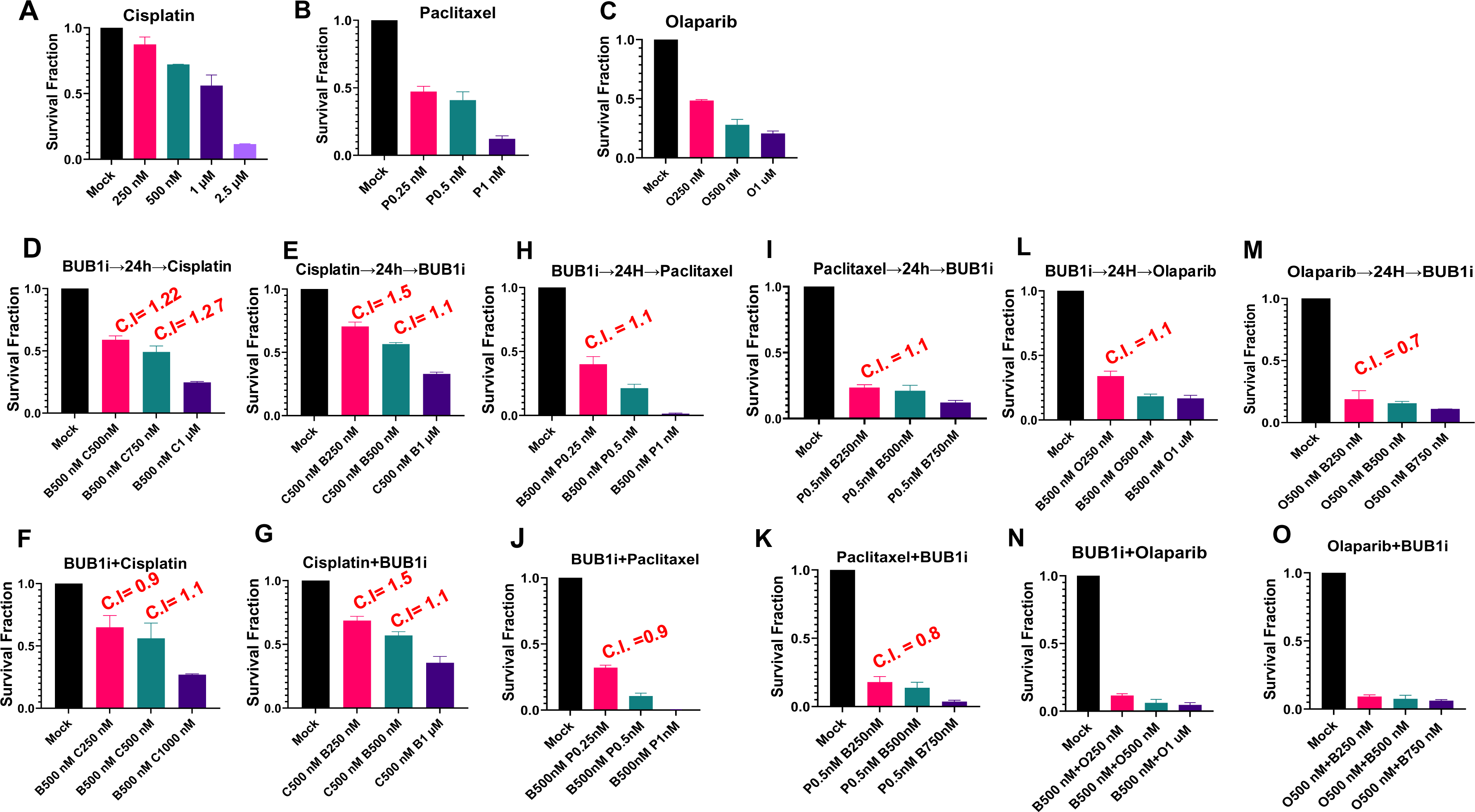
Clonogenic assay demonstrating effect of BUB1i on chemotherapy efficacy in H1975. (A-C) Drug alone (D-I) Sequential treatment (J-O) Concurrent treatment of drugs. C.I. >1, = 1 and <1 indicate antagonism, additivity, and synergism, respectively.

### 3.5. BUB1 inhibition chemosensitizes NSCLC in double and triple drug combinations

BUB1i was antagonistic with cisplatin in A549 and H2030 (**Figures 5A-B**) with additivity seen at higher drug doses in A549 (**Figure 5A**). BAY1816032 was synergistic with olaparib in A549 (**Figure 5C)** while it was synergistic or additive in H2030 depending on the doses (**Figure 5D**). BAY1816032 enhanced cell killing when combined with paclitaxel in NSCLC, showing synergy (C.I. < 1) (**Figures 5E-F**). Combining BAY1816032 with paclitaxel and olaparib (triple drug) further enhances cell killing in A549, H2030 and H1975 (**Figures 5G-I**), suggesting potential therapeutic benefits in NSCLC.

**Figure 5:**
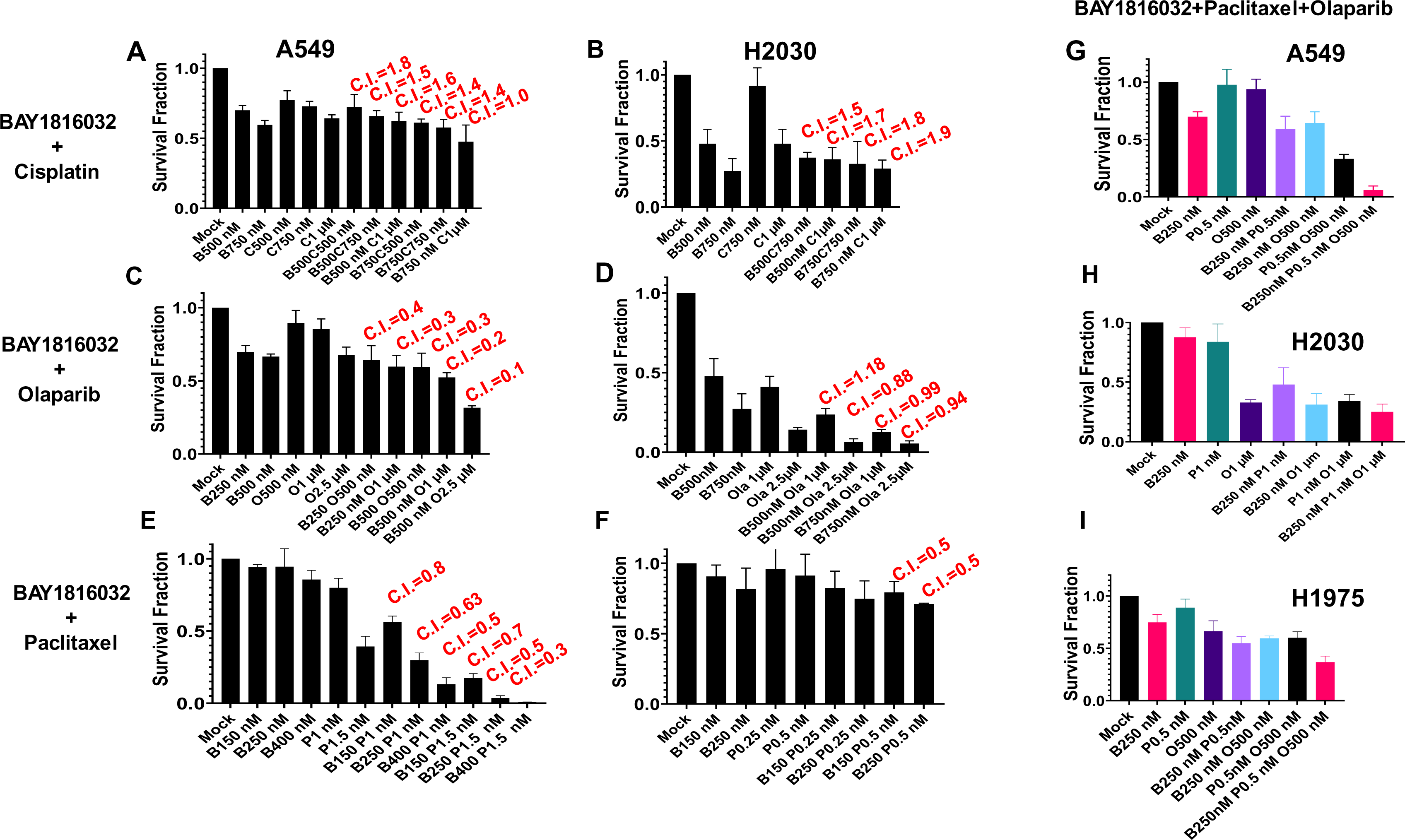
BUB1i causes chemo-sensitization with olaparib and paclitaxel in NSCLC. Double drug combination: BUB1i showed antagonism with cisplatin in (A) A549 and (B) H2030 with additivity seen at higher drug doses in A549. BAY1816032 demonstrated synergism with olaparib in A549 (C) while it was synergistic or additive in H2030 depending on the doses (D). BAY1816032 enhanced cell killing when combined with paclitaxel in NSCLC, showing synergy (C.I. < 1) (E-F). Combining BAY1816032 with paclitaxel and olaparib (triple drug) further enhances cell killing in A549, H2030 and H1975 (Figures 5G-I), suggesting potential therapeutic benefits in NSCLC.

### 3.6. BUB1 inhibition sensitizes NSCLC to radiation

BAY1816032 demonstrated enhanced radiation-induced cell killing in A549, H2030, and H1975 cells, with the radiation enhancement ratio (rER) greater than 1 (**Figures 6A-C)**. Higher doses of BAY1816032 corresponded to increased rER values. In Calu 1 cells, the IC_50_ for BAY1816032 alone was 4µM, (**Figure 2E**) which decreased to 2µM when combined with radiation (4Gy) (**Supplementary Figure 1A**). Similarly, the IC_50_ for paclitaxel, cisplatin and Olaparib alone also decreased from 7.9nM, 4.6µM and >15µM (**Figures 4E, 4L and 4S**) to 5.5nM, 2µM and 8.5µM respectively when combined with radiation (4Gy) (**Supplementary Figures1B-D**). However, in normal lung epithelial PCS-300-010 cells, BAY1816032 did not exhibit radiation sensitization, with an IC_50_ >15µM both alone (**Figure 2A**) and with radiation (**Supplementary Figure 1E**). Further, IC_50_ for paclitaxel, cisplatin and Olaparib alone (**Figures 4A, 4H and 4O**) did not change when combined with radiation in PCS-300-010 cells (**Supplementary Figure 1F-H**). These results indicate the potential of BAY1816032 as a radiosensitizer for NSCLC while potentially minimizing toxicity in normal tissues.

**Figure 6:**
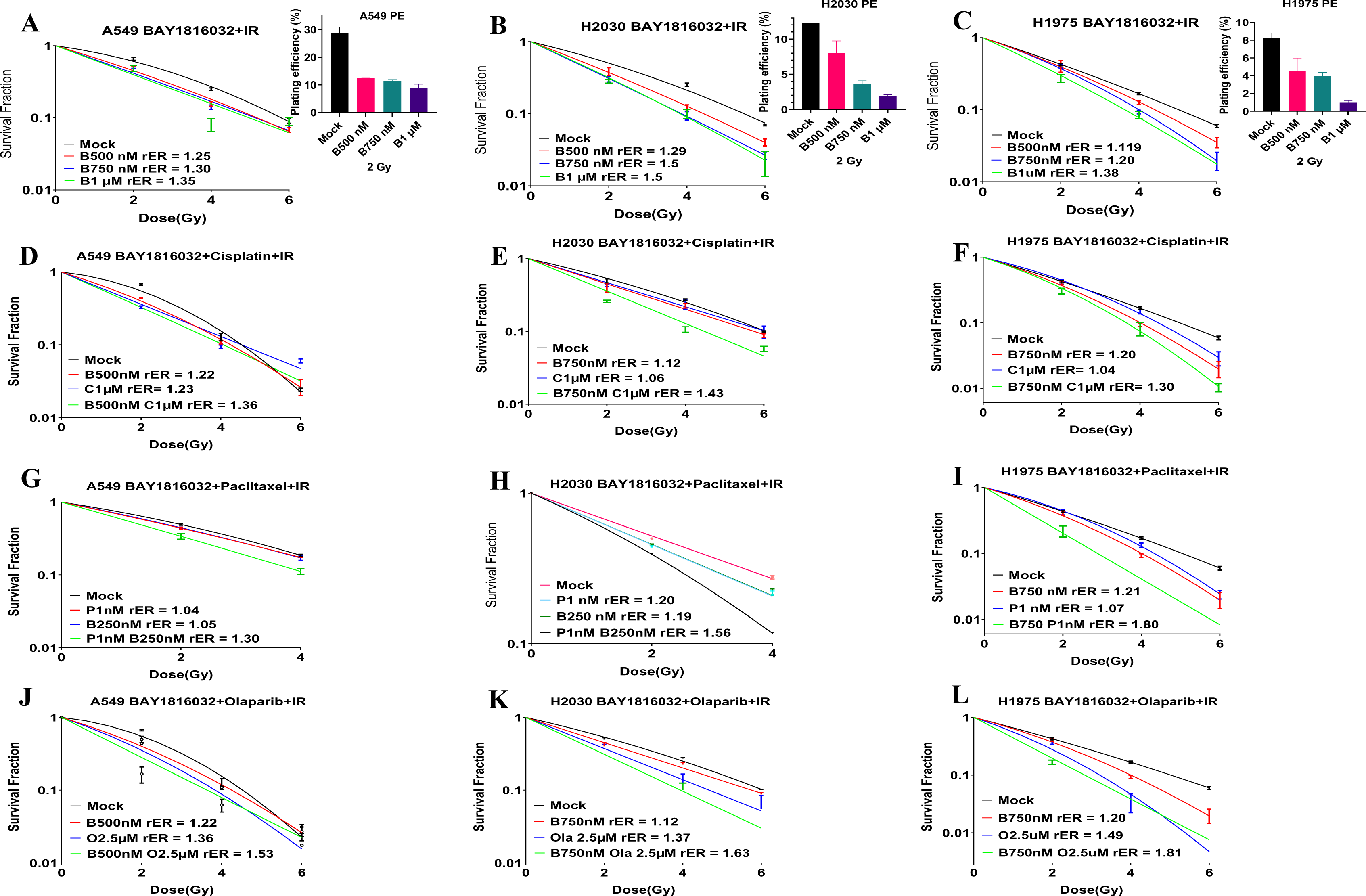
BUB1 inhibition radiosensitizes NSCLC (A) A549 (B) H2030 (C) H1975. BAY1816032 leads to chemo-radiosensitization in NSCLC: combination treatment of BAY1816032 and another drug with radiation lead to increased sensitization to show synergistic effects. BAY1816032+cisplatin+IR in (D) A549 (E) H2030 (F) H1975, BAY1816032+paclitaxel+IR in (G) A549 (H) H2030 (I) H1975, BAY1816032+olaparib+IR in (J) A549 (K) H2030 (L) H1975.

### 3.7. BUB1 inhibition sensitizes NSCLC and SCLC to chemoradiation

BAY1816032 led to strong chemoradiation sensitization in NSCLC with platinum (BAY1816032+cisplatin+IR; **Figures 6D-F**), anti-microtubular agent (BAY1816032+paclitaxel+IR; **Figures 6G-I**), and with PARP inhibitor (BAY1816032+olaparib+IR; **Figures 6J-L**). This potential synergistic sensitization increased with increasing radiation doses across all tested combinations (**Figures 6D-L**).

The response of SCLC to BUB1i was variable in double drug or radiation combinations. BUB1i led to synergistic sensitization with paclitaxel in NCI-H1876 and NCI-H2198 which was independent of radiation (**Figures7A-B**). Although the effect of BUB1i with cisplatin in NCI-H1876 cell line was antagonistic (**Figure 7C**; black bars), it shifted to synergism/additivity when combined with radiation (**Figure 7C**; magenta bars). BUB1i demonstrated synergistic sensitization with cisplatin in NCI-H2198 cells (**Figure 7D**) which was independent of radiotherapy. BAY1816032 combination with olaparib was synergistic in both the SCLC cell lines (**Figures 7E-F**).

**Figure 7:**
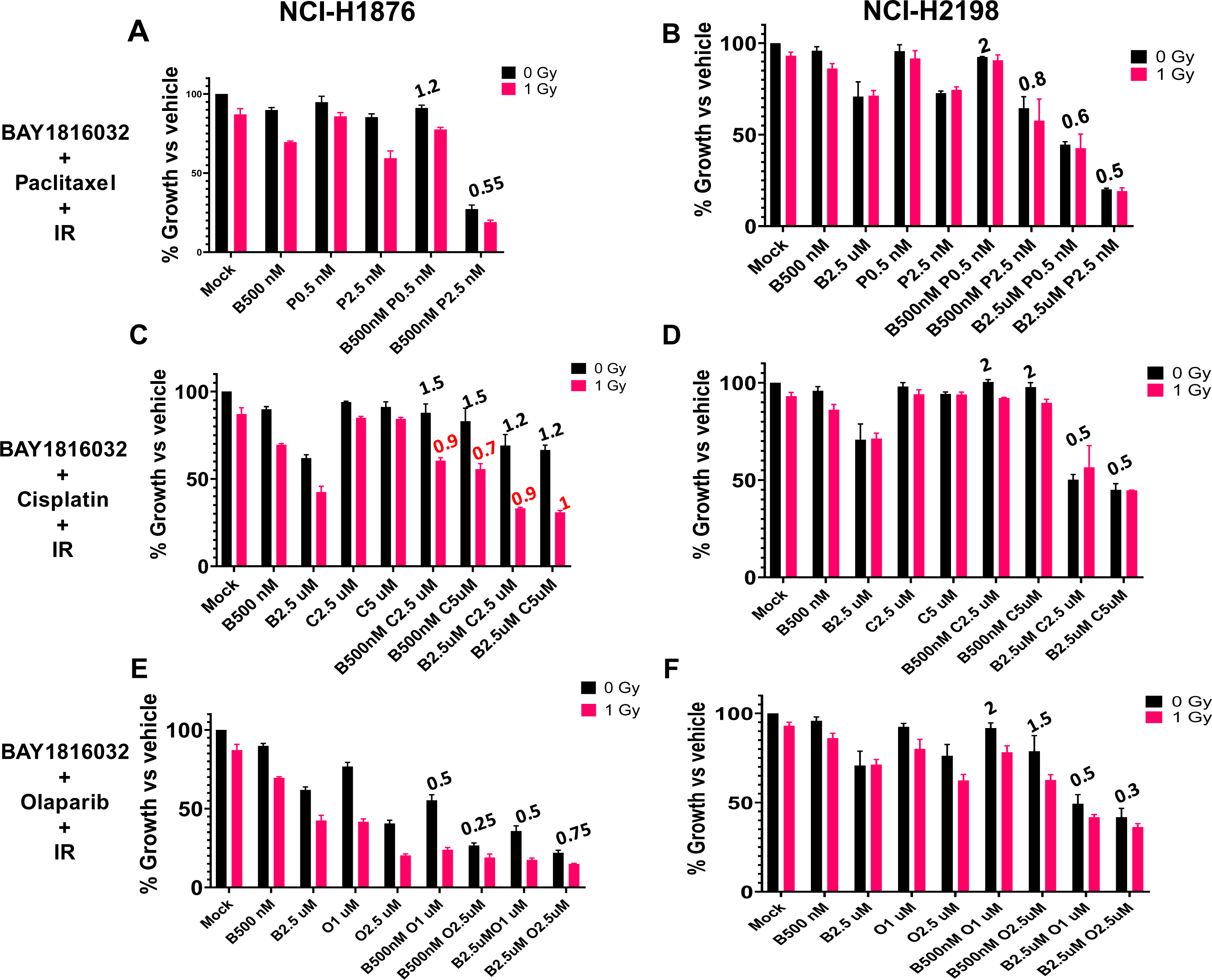
BUB1i causes chemo-radiosensitization with olaparib and paclitaxel in SCLC in Alamar blue assay with or without radiation. BAY181062 demonstrated synergistic sensitization with paclitaxel in (A) NCI-H1876 and (B) NCI-H2198, independent of radiation. BAY181062 with cisplatin in NCI-H1876 cell line was antagonistic (C; black bars), it shifted to synergism/additivity when combined with radiation (C; magenta bars). BAY181062 demonstrated synergistic sensitization with cisplatin in NCI-H2198 cells (D), independent of radiotherapy. BAY1816032 combination with olaparib was synergistic in both the SCLC cell lines (E-F).

### 3.8. BUB1 inhibition delays DNA double strand break repair and elicits pro-apoptotic and anti-proliferative responses

In H2030 cells, BUB1 inhibition led to persistence of γ-H2AX foci 24 hours post-irradiation **(Figures 8A-B**) while control cells returned to baseline levels indicating that DNA double-strand break repair pathway was impacted. BAY1816032 with radiation altered key cellular markers associated with apoptosis and proliferation. Increased BAX and decreased BCL2 expression shifted the cellular balance toward pro-apoptotic signaling (**Figure 8C**). PCNA expression decreased, indicating downregulation of cellular proliferation. Elevated expression of Caspase 9, Caspase 3 and TP53BP1 (**Figure 8C**) indicate enhanced apoptotic response post treatment.

**Figure 8:**
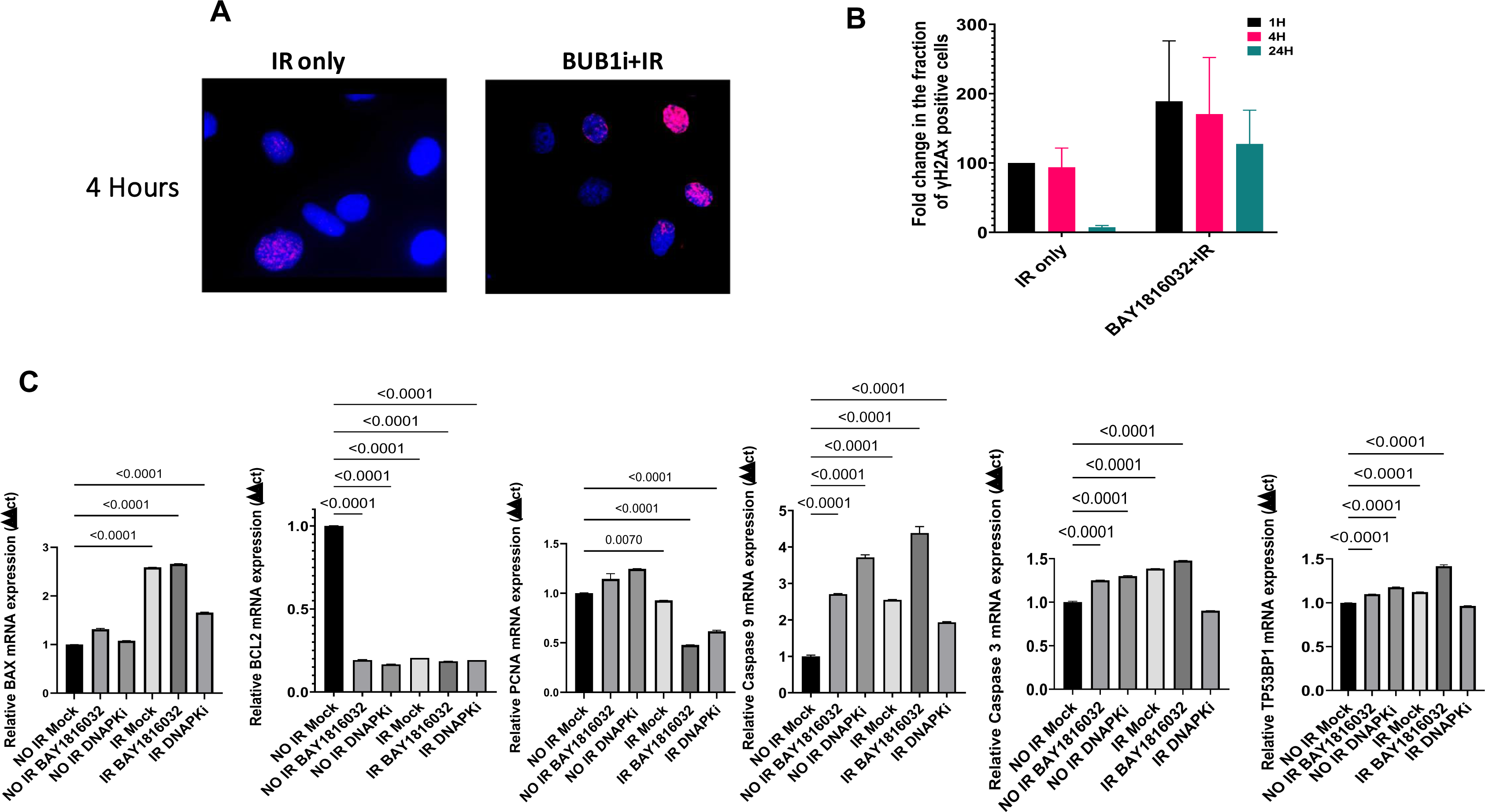
BUB1i delays DNA double-strand break repair. (A) BUB1i inhibition (BAY1816032) causes delay in H2AX foci resolution in H2030. (B) γH2AX positive foci quantification. (C) qPCR: BUB1 inhibition causes increase in pro-apoptotic markers in A549.

## 4. Discussion

The observation that BUB1 is upregulated across various lung cancer subtypes (**Figures 1A-D**) emphasizes its potential as a crucial molecular target in lung cancer. Our analyses are consistent with previous studies that have reported BUB1 upregulation in solid tumors (11–13, 21). BUB1’s association with TP53 mutations in LUAD and LUSC (**Figures 1H-1I**) suggests involvement in the p53 signaling pathway (22). This is consistent with its recognition as a potential synthetic lethal gene with p53 (23). Wang *et. al.* (2020) established that high BUB1 levels correlate with poorer survival and multiple oncogenic signaling pathways (22). In our analyses, the correlation of higher BUB1 expression with poorer survival was also observed (**Figure 1E**). Confirmation of BUB1 overexpression in tumor tissues (**Figures 1H-I and 1L**) and lung cancer cell lines (**Figure 1M**) further emphasizes its significance in lung cancer. No correlation between BUB1 expression and cancer stage or grade was observed (**Figures 1J-K**), consistent with findings in other cancers including LUAD (12, 24).

BUB1i with BAY1816032 resulted in a significant dose-dependent reduction in proliferation in both NSCLC and SCLC (**Figures 2B-G**). Normal lung cells showed toxicity at ∼4 times the dose required for cancer cells (**Figure 2A**), indicating the specificity in targeting cancer cells by BAY1816032. Clonogenic assay further validated the lasting impact of BAY1816032 on reducing clonogenic capacity, at much lower doses (**Figures 2I-K**). Chemotherapy drugs showed IC_50_ at micro or nanomolar range in the cell lines studied (**Figures 3A-U**). Concurrent drug treatment was found to be the best approach for drug combination (**Figures 4J-O**). While BAY1816032’s efficacy in double drug combinations have been established previously elsewhere (17), our study additionally confirmed BAY1816032 causes chemo-sensitization in triple drug combinations (**Figures 5G-I**). Further, we present the novel finding that BAY1816032 chemo-sensitizes SCLC (**Figures 7A-H**).

BUB1 has been recognized as a potential target in radiation sensitization (25), and is reported to play an integral role in DNA damage response to radiation (9, 26–29). Our results established BAY1816032 radiosensitizes NSCLC (**Figures 6A-C**). Triple combinations involving BUB1i, chemotherapy and radiation causes chemo-radiosensitization in both NSCLC and SCLC (30, 31) (**Figure 6D-L** and **Figure 7A-F**). BUB1 inhibition resulted in the persistence of γ-H2AX foci in H2030 (**Figure 8A** and **8B**), strongly indicating a delayed DNA damage repair in the absence of BUB1 kinase activity. Increased levels of 53BP1 in response to BAY1816032 treatment and radiation suggest an active DNA damage response (32), with BUB1 inhibition impairing repair kinetics. Altered expression of key markers such as BAX, BCL2, PCNA, and Caspases 9 and 3 (**Figure 8C**) following BAY1816032 and radiation indicates a shift toward a pro-apoptotic and anti-proliferative state. Similar pro-apoptotic responses have been reported in other cancers in response to BUB1 inhibition (16, 33), emphasizing a potential commonality in the molecular responses triggered by BUB1 inhibition across different cancer models.

PARP1/2 proteins usually detect SSB and recruit factors to repair the SSB. PARPi causes either PARP trapping on DNA break sites that lead to replication fork collapse and cell death (specially in BRCA mutant cell lines (34)) or PARPi can upregulate NHEJ and reduce HR leading to genomic instability and cell killing (35). We anticipate that BUB1i sensitizes to PARP inhibitors because PARPi can increase the dependency on NHEJ for which BUB1 is critical. Platinums (cisplatin, carboplatin) form DNA adducts that causes DNA replication errors leading from SSB to DSB, cell-cycle arrest in G1-S and ultimately cell death (36, 37). Increased platinum-DNA adduct repair has been shown to be associated with Cisplatin resistance (38). Similarly, Paclitaxel blocks depolarization of microtubules, leading to improper chromosome segregation, G2/M cell-cycle arrest (39, 40) resulting in apoptotic cell-death (41, 42). NSCLC cell lines have variable sensitivity to these drugs. BUB1 not only regulates cell-cycle but it also regulates DNA damage response. Therefore, we propose that BUB1 inhibition sensitizes to these agents and overcome resistant because of its ability to target multiple pathways. In summary, our data demonstrate that BUB1i increases NSCLC cell death by Paclitaxel, Cisplatin and Olaparib (in BRCA efficient cells) and that these effects are accentuated when combined with radiation. The observed sensitization effects of BUB1 inhibition in lung cancer suggest complex interactions involving DNA damage repair pathways and modulation of apoptotic signaling. Future studies will decipher these complex mechanisms.

## 5. Conclusion

This study demonstrated BUB1 to be a promising therapeutic target in lung cancer. Inhibiting BUB1 enhanced the sensitivity of both NSCLC and SCLC to a variety of chemotherapies including cisplatin and paclitaxel, targeted therapy (PARPi) and radiation. These findings highlight the potential of BUB1 inhibition as a promising approach to enhance the effectiveness of radiotherapy and chemoradiation in treating lung cancers.

## Acknowledgements

The authors thank Histology core-HFH for immunohistological staining. Grant support to SN (NIH/NCI R21 CA252010, Henry Ford Cancer Institute (HFCI) and Henry Ford Health Research Administration Start Up grant, HFH Proposal Development Award, HFH Near Miss Award, HFH Radiation Oncology Start Up grant.

## Data availability

Data available from the corresponding author on reasonable request.

## Author contributions

SN; Experiments: SS, ST

## Consent for publication

Not applicable

---

**Supplementary Figure 1:** (A-D) Radiation enhanced cell killing when combined with drugs in Calu-1. (F-H) No effect of radiation on drug only treatment in PCS-300-010 normal lung epithelial cells.

