## Supplementary figures and images for "BUB1 inhibition sensitizes lung cancer cell lines to radiotherapy and chemoradiotherapy"

### Supplementary Figure 1

#### Slide 1
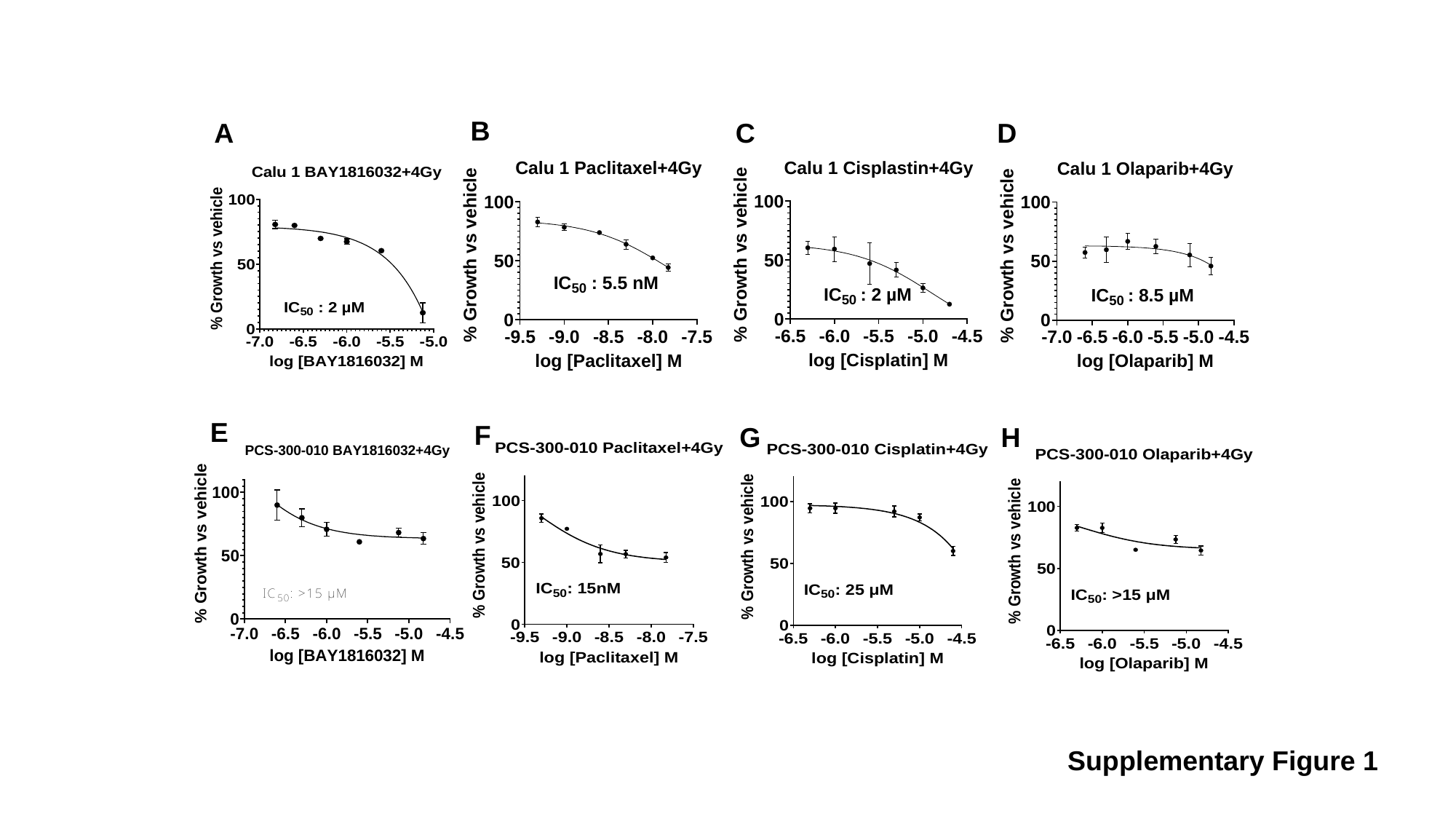

B
A
C
D
E
F
G
H
Supplementary Figure 1
